# The P132H mutation in the main protease of Omicron SARS-CoV-2 decreases thermal stability without compromising catalysis or small-molecule drug inhibition

**DOI:** 10.1101/2022.01.26.477774

**Authors:** Michael D Sacco, Yanmei Hu, Maura V Gongora, Flora Meilleur, M Trent Kemp, Xiujun Zhang, Jun Wang, Yu Chen

**Affiliations:** Department of Molecular Medicine, Morsani College of Medicine, University of South Florida, Tampa, FL, 33612, United States; Department of Medicinal Chemistry, Ernest Mario School of Pharmacy, Rutgers, the State University of New Jersey, Piscataway, NJ, 08854, United States; Neutron Scattering Division, Oak Ridge National Laboratory, Oak Ridge, TN, 37830, United States

**Author notes:** Corresponding authors: Yu Chen, Jun Wang.

## Abstract

The ongoing SARS-CoV-2 pandemic continues to be a significant threat to global health. First reported in November 2021, the Omicron variant (B.1.1.529) is more transmissible and can evade immunity better than previous SARS-CoV-2 variants, fueling an unprecedented surge in cases. To produce functional proteins from this polyprotein, SARS-CoV-2 relies on the cysteine proteases Nsp3/papain-like protease (PLpro) and Nsp5/Main Protease (M^pro^)/3C-like protease to cleave at three and more than 11 sites, respectively.^1^ Therefore, M^pro^ and PL^pro^ inhibitors are considered to be some of the most promising SARS-CoV-2 antivirals. On December 22, 2021, the Food and Drug Administration (FDA) issued an Emergency Use Authorization (EUA) for PAXLOVID, a ritonavir-boosted formulation of nirmatrelvir. Nirmatrelvir is a first-in-class orally bioavailable SARS-CoV-2 M^pro^ inhibitor.^2^ Thus, the scientific community must vigilantly monitor potential mechanisms of drug resistance, especially because SARS-CoV-2 is naïve to M^pro^ inhibitors. Mutations have been well identified in variants to this point.^3^ Notably, Omicron M^pro^ (OM^pro^) harbors a single mutation– P132H. In this study we characterize the enzymatic activity, drug inhibition, and structure of OM^pro^ while evaluating the past and future implications of M^pro^ mutations.

## RESULTS AND DISCUSSION

Using an established FRET assay to assay proteolytic activity, we show OM^pro^ and WT M^pro^ have equivalent affinity and catalytic constants for its substrate (**Fig 1A**). However, subsequent melting temperature experiments reveal OM^pro^ has a T_m_ of 53.6 ± 0.1 °C; lower than the WT M^pro^ T_m_ of 56.2 ± 0.2 by 2.6 °C (**Fig. 1B**). Despite its lower melting temperature, OM^pro^ and WT M^pro^ hydrolyze their substrates at comparable rates for up to 25 hours at 37°C (**Fig. 1C**). Further biochemical analysis suggests that OM^pro^ and M^pro^ are equally susceptible to covalent inhibitors such as GC-376, PF-07321332 (nirmatrelvir), and PF-00835231 (**Fig. 1D**). Interestingly, when these inhibitors are incubated with WT M^pro^ and OM^pro^ their melting temperatures become nearly identical, despite the different melting temperature of their apo forms (**Fig. 1E**). This suggests that inhibitors may actually stabilize OM^pro^ to a greater extent than WT M^pro^. These described biochemical/biophysical results are in line with the equivalent antiviral potency of these molecules against WT and Omicron and the lack of observable differences in the active site.^4-7^

**Fig. 1.**
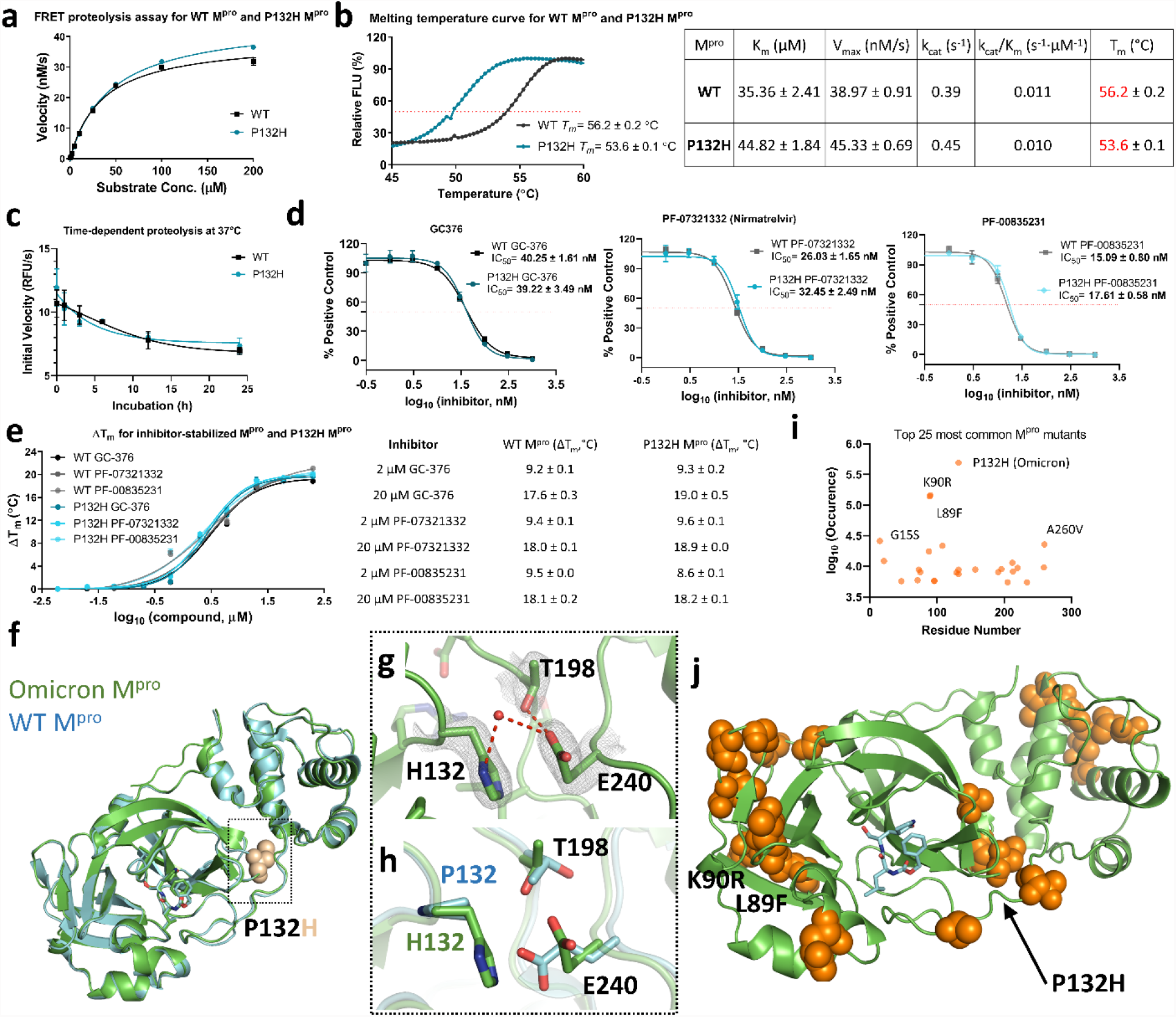
Biochemical and structural comparison of WT and Omicron M^pro^ P132H. (A) Characterization of enzymatic activity. (B) Thermal shift assay for apo proteins. (C) Time dependent proteolytic activity. (D) Inhibition by covalent inhibitors GC-376, PF-07321332 (nirmatrelvir), and PF-00835231. (E) Thermal shift assay for GC-376, PF-07321332 (nirmatrelvir), and PF-00835231. (F) Crystal structure of SARS-CoV-2 Omicron M^pro^ + GC-376 at 2.05 Å resolution superimposed with WT M^pro^ + GC-376 (blue; PDB ID 7C6U). P132H is shown as spheres. (G) Electron density map of H132 and surrounding residues. 2F_o_-F_c_ map is contoured at 1 s and shown in grey. (H) Structural comparison of position 132 and interacting residues in WT M^pro^ and Omicron M^pro^. (I) Top 25 most common M^pro^ mutants at each position. (J) Top 25 most common M^pro^ mutants mapped onto the crystal structure of SARS-CoV-2 OM^pro^.

In parallel to our study, Ullrich *et al*. also characterized six M^pro^ mutants identified from the circulating SARS-CoV-2 variants.^8^ The k_cat_/K_m_ values for WT and Omicron M^pro^ were 0.016 and 0.023 S^-1^µM^-1^, which were similar to the values obtained in our study (**Fig. 1B**). In addition, the Pfizer team reported PF-07321332 (nirmatrelvir) retained potent inhibition against Omicron P132H M^pro^ (K_i_ = 0.635 nM), similar to the potency of the WT (K_i_ = 0.933 nM).^9^ Our independent study further confirmed these results.

Crystals of OM^pro^ in complex with GC-376 were collected at 2.05 Å resolution in the I 1 2 1 space group with a R_work_/R_free_ of 0.179 / 0.219 (Supplementary information, Table S1; PDB ID 7TOB). The unit cell is the same as many previously solved WT M^pro^ crystal structures where a= 45.19 Å, b= 52.99 Å, c= 113.01 Å and α =90.00°, β= 100.50°, γ= 90.00°. Overall, OM^pro^ has a nearly identical structure to WT M^pro^ (**Fig. 1F**).^10^ The most pronounced differences involve the area around the mutation associated with OM^pro^, P132H. Found 22 Å from the catalytic cysteine Cys145, P132H lies between the catalytic domain and the dimerization domain, thus, it does not impart any direct structural changes to the active site.

Clear electron density shows that His132 forms π-stacking interactions with the sidechain of Glu240 (**Fig. 1G**). As the imidazole sidechain of histidine is protonated near physiological pH, the interaction with Glu240 may be further strengthened through electrostatic interactions. Additionally, there is a newly formed water mediated hydrogen bond with Glu240. We also find that Glu240 reorients itself towards the core to accommodate the bulkier His sidechain, where it forms a new hydrogen bond with Thr198 (**Fig. 1H**). Consequently, the Thr198 sidechain rotates ∼ 90°, placing its hydroxyl group in close distance to Glu240. As a result of these conformational changes, certain portions of the enzyme appear to move very slightly, ∼ 0.5 Å (**Fig. 1F**). While the overall structure of M^pro^ and OM^pro^ align with an RMSD of 0.413 Å, the catalytic domains (1-184) and dimerization domain (185-306) can be individually aligned with RMSD values of 0.337 Å and 0.309 Å, suggesting they move away from the other. We speculate that the lower thermal stability of apo OM^pro^ may be due to minor residue adjustments to accommodate the bulkier His132 sidechain, ultimately destabilizing its structure. Because the mutation occurs at the interface between the dimerization domain and the catalytic domain, these movements can also affect intramolecular packing. Residue 132 is also located at the turn between two β-sheets, a position that naturally favors the cyclic sidechain of Pro.

Although the P132H mutation does not appear to reduce enzymatic activity and inhibitor binding (**Fig. 1 A-D**), the decrease in thermal stability (**Fig. 1B**) indicates protein flexibility may be greater, which plays an important role in enzyme evolution, especially to broaden substrate profile or alter ligand binding. M^pro^ can recognize a wide range of peptide substrates, although its P1 position preferentially binds to glutamine. Future studies investigating whether P132H and other mutations can influence enzymatic activity for larger substrate libraries or other known ligands will prove useful.

Extensive sequencing of SARS-CoV-2 isolates has provided unprecedented insights to the mutations that occur in the viral RNA genome.^3^ Based on the annotations provided through CoVsurver enabled by GISAID (www.gisaid.org/epiflu-applications/covsurver-mutations-app), mutations of NSP5 appear mostly stochastic (Supplementary information, Figure 1), however, several positions have a disproportionately large number of mutations. Of the top 25 most common mutants (**Fig. 1I**), three are found on P132: P132H (489,444 occurrences; EPI_ISL_8931050), P132L (8,813 occurrences; EPI_ISL_8768027), and P132S (7,452 occurrences; EPI_ISL_8925342). L89F, K90R, and K88R are the second, third, and seventh, most common mutations, suggesting this β-sheet is also a hotspot. Notably, none of the 25 most common mutants involve residues in the active site or at the dimerization interface (**Fig. 1J**). With several exceptions, including P132H, the resulting amino acid is often similar in size and physicochemical properties, such as K→ R. However, SARS-CoV-2 has not yet encountered M^pro^ antivirals. If nothing else, SARS-CoV-2 has taught us that widespread proliferation, low fidelity genome synthesis, and selective pressure will quickly produce drug-resistant phenotypes. Thus, it is important to monitor future mutations and their associated biochemical properties to anticipate future drug-resistance.

## DATA AVAILABILITY

The X-ray crystal structure of the Omicron M^pro^ P132H mutant in complex with GC376 was deposited in PDB with the code 7TOB.

## ACKNOWLEDGEMENTS

This research used resources at the X-ray home source operated by the Oak Ridge National Laboratory, a DOE Office of Science User Facility. This research was supported by the National Institutes of Health (NIH) (grants AI147325, AI157046, and AI158775) to J.W. We thank Eric M Lewandowski for reviewing this manuscript.

## AUTHOR CONTRIBUTIONS

M.D.S., J.W., and Y.C. conceived the experiments, interpreted the data, and composed the manuscript. Y.H. performed the biochemical assays and melting temperature experiments. Proteins were cloned and purified by X.J. Crystal structures were solved and refined by M.D.S. Crystals were grown by M.V.G. X-ray datasets were prepared and collected by M.T.K and F.M.

## COMPETING INTERESTS

The authors declare no competing interests

## ADDITIONAL INFORMATION

## Supplementary information

The online version contains supplementary material available at

## Correspondence

and requests for materials should be addressed to Jun Wang or Yu Chen.

## Notes

### Competing Interest Statement

The authors have declared no competing interest.

