## Supplementary Data for "The P132H mutation in the main protease of Omicron SARS-CoV-2 decreases thermal stability without compromising catalysis or small-molecule drug inhibition"

*Corresponding authors:

**Materials and Methods**

**Protein purification and crystallization**
Using QuikChange site directed mutagenesis, Pro132 was mutated to His in a pET29a(+) vector with the cloned gene for the BetaCoV/Wuhan/WIV04/2019 SARS CoV-2 main protease, producing a gene that is equivalent to that found in the Omicron SARS-CoV-2. Protein was then expressed and purified as previously described^1^. Crystals of OM^pro^ with GC-376 were grown by incubating 15 mg/mL OM^pro^ overnight at 4 °C with two-fold molar excess GC-376. Protein was diluted to 5 mg/mL in gel filtration column buffer (20 mM Tris pH 8.0, 150 mM NaCl) and DMSO was added for a final concentration of 4 %. OM^Pro^ was then mixed with crystallization buffer (20% PEG 3350, 0.2 M KNO_3_) and allowed to grow in a hanging drop apparatus at 20 °C for approximately two weeks.

**X-ray dataset collection**
A plate-like crystal with sides of ~ 200 x 150 µm was transferred to a cryo-protectant solution of 27.5 % PEG 3350, 0.2 M KNO_3_, and 15% glycerol, flash frozen in liquid nitrogen and transferred to an Oxford Cryosystems nitrogen cryostream for data collection. An X-ray data set was collected at 100 K on a Rigaku MicroMax-007 HF microfocus rotating anode X-ray generator equipped with a Dectris EIGER R 4M detector. A total of 180 frames were collected with an exposure time of 120 s per frame and Δφ of 1°. The data were indexed and integrated using *CrysAlis^Pro^* (Rigaku, The Woodlands, Texas, USA) and scaled and merged with AIMLESS^2^ in the CCP4 suite. The final model was solved via molecular replacement using PDB ID 7C6U as the reference model for Molrep^3^. This structure was subsequently refined using Refmac5^4^ and Coot^5^.

**Mutation analysis**
Sequence data for SARS-CoV-2 isolates was gathered using the CoVsurver of the Global Initiative on Sharing Avian Influenza Data, Developed by A*STAR Bioinformatics Institute (BII), Singapore. All numbers shown are accurate as of Jan 15, 2022.

### Enzymatic assays For determination of *K*_m_ and *V*_max_, WT or P132H M^pro^ was diluted in the reaction buffer (20 mM HEPES (pH 6.5), 120 mM NaCl, 0.4 mM EDTA, 4 mM DTT, and 20% glycerol) to the final concentration of 100 nM, and various concentrations (0, 0.1, 0.25, 0.5, 1, 2.5, 5, 10, 25, 50, 100, 200 µM) of M^pro^ specific FRET substrate were added to initiate the reaction^1,6^. The reaction was monitored every 90 seconds for 1 hour at 30 °C in BioTek Cytation 5 (Agilent, Santa Clara, CA) with excitation wavelength at 360/40 nm and emission wavelength at 460/40 nm. The initial velocity of the enzymatic reaction at each concentration of FRET substrate was determined in the first 15 min by linear regression. The *K*_m_ and *V*_max_ were determined by fitting the curves with nonlinear regression of initial velocity vs. concentration of FRET substrate using the Michaelis-Menten equation in Prism 8.

IC_50_ values of M^pro^ inhibitors, 100 nM WT or P132H M^pro^ were determined by incubating serial concentrations (0, 0.001, 0.003, 0.01, 0.03, 0.1, 0.3, 1 µM) of inhibitors at 30 °C for 30 min in the reaction buffer. The reaction was initiated by adding 10 µM FRET substrate and monitored every 90 seconds for 1 h at 30 °C, and the initial velocity was determined in the first 15 min by linear regression. The IC_50_ values were determined by fitting the curves with nonlinear regression using log (concentration of inhibitor) vs. response with variable slopes in Prism 8.

Time-dependent proteolysis was performed by diluting WT or P132H M^pro^ to 100 nM in the reaction buffer and incubating at 37 °C for 0, 1, 3, 6, 12, 24, or 48 hrs. At each time point, 10 µM FRET substrate was added to initiate the reaction, which was monitored every 90 seconds for 1 h at 30 °C. Initial velocity was determined by linear regression for the first 15 min.

### Differential scanning fluorimetry The binding of M^pro^ inhibitors to WT or P132H M^pro^ was determined by differential scanning fluorimetry (DSF) using QuantStudio^TM^ 5 Real-Time PCR System (Thermo Fisher Scientific, [Waltham, MA](https://www.google.com/search?sxsrf=AOaemvJBwxgyAScKwNKSQGdW3rfXiTuSfg:1643003646962&q=Waltham&stick=H4sIAAAAAAAAAOPgE-LSz9U3MCooMTBJU-IAsTOqjE21tLKTrfTzi9IT8zKrEksy8_NQOFYZqYkphaWJRSWpRcWLWNnDE3NKMhJzd7AyAgDThZNCUQAAAA&sa=X&ved=2ahUKEwiwlffF2cn1AhULjIkEHenSDdcQmxMoAXoECCoQAw)), as previously described^7,8^. DSF was carried out in a 96-well PCR plate by mixing 3 μM of WT or P132H M^pro^ protein with various inhibitor concentrations (0, 0.02, 0.06, 0.2, 0.6, 2, 6, 20, 60, 200 µM), and incubating at 30 °C for 30 min. 1× SYPRO orange (Thermo Fisher Scientific, [Waltham, MA](https://www.google.com/search?sxsrf=AOaemvJBwxgyAScKwNKSQGdW3rfXiTuSfg:1643003646962&q=Waltham&stick=H4sIAAAAAAAAAOPgE-LSz9U3MCooMTBJU-IAsTOqjE21tLKTrfTzi9IT8zKrEksy8_NQOFYZqYkphaWJRSWpRcWLWNnDE3NKMhJzd7AyAgDThZNCUQAAAA&sa=X&ved=2ahUKEwiwlffF2cn1AhULjIkEHenSDdcQmxMoAXoECCoQAw)) was added into each well and the fluorescence was monitored under a temperature gradient ranging from 20 to 95 °C (incremental steps of 0.05 °C/s). The melting temperature (*T_m_*) was calculated as the mid-log of the transition phase from the native to the denatured protein using the Boltzmann model in Protein Thermal Shift Software v1.3. The melting temperature shift (Δ*T_m_*) was calculated by subtracting *T_m_* of protein in the presence of DMSO from *T_m_* in the presence of inhibitors.

**Table S1. Crystallization statistics for Omicron SARS-CoV-2 M^pro^.**

| PDB ID | 7TOB |
| --- | --- |
| Protein | Omicron SARS-CoV-2 M^pro^ |
| Ligand | GC376 |
| **Data Collection** |  |
| Space Group | I 1 2 1 |
| Cell Dimensions |  |
| a, b, c (Å) | 45.19, 52.99, 113.01 |
| α, β, γ (°) | 90.00, 100.50, 90.00 |
| Resolution (Å) | 26.20 – 2.05 |
|  | (2.11 – 2.05) |
| R_merge_ | 0.065 (0.462) |
| <I>/σ<I> | 9.4 (2.2) |
| Completeness (%) | 99.2 (98.4) |
| Redundancy | 3.3 (2.9) |
| **Refinement** |  |
| Resolution (Å) | 26.20 – 2.05 |
|  | (2.13 – 2.05) |
| No. reflections/free | 16,426 / 1,606 |
| R_work_/R_free_ | 0.179 / 0.219 |
| Clashscore | 4.42 |
| No. Atoms |  |
| Overall | 2,613 |
| Protein | 2,372 |
| Ligand/Ion | 43 |
| Water | 198 |
| B-Factors (Å^2^) |  |
| Overall | 32.84 |
| Protein | 32.20 |
| Ligand/Ion | 37.77 |
| Solvent | 39.35 |
| RMS Deviations |  |
| Bond Lengths (Å) | 0.014 |
| Bond Angles (°) | 1.87 |
| Ramachandran Favored (%) | 97.37 |
| Ramachandran Allowed (%) | 2.63 |
| Ramachandran Outliers (%) | 0.00 |
| Rotameric  Outliers (%) | 1.14 |

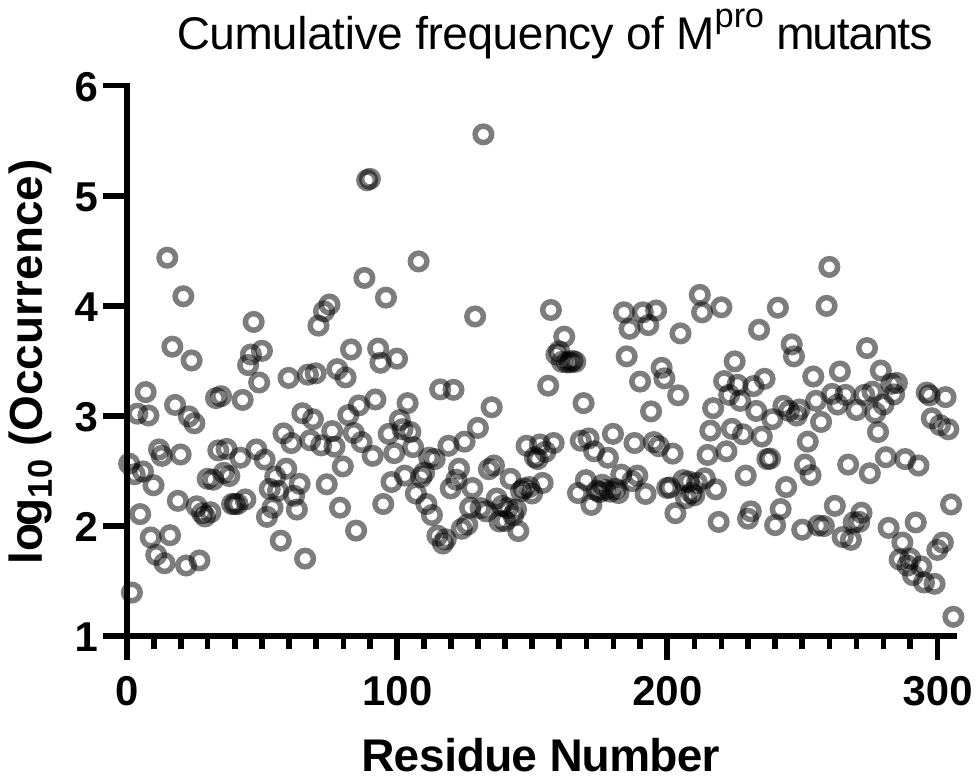
Figure S1. Cumulative frequency of M^pro^ mutants at each position
